# Composition, growth, succession, and function in the *Cladophora* microbiome: insights from quantitative Stable Isotope Probing and NanoSIMS imaging

**DOI:** 10.1101/2025.05.12.652736

**Authors:** Raina Fitzpatrick, Bruce Hungate, Mary Power, Megan Foley, Ty Samo, Egbert Schwartz, Micheala Hayer, Peter K. Weber, Jennifer Pett-Ridge, Jane Marks

**Affiliations:** Department of Biological Sciences and Center for Ecosystem Science and Society, Northern Arizona University, Flagstaff AZ 86011; Department of Integrative Biology, University of California, Berkeley CA 94720; Physical and Life Sciences Directorate, Lawrence Livermore National Lab, Livermore, California, USA; Life & Environmental Sciences Department, University of California Merced, CA, USA; Innovative Genomics Institute, University of California Berkeley, Berkeley CA, USA

## Abstract

The branching green macroalga *Cladophora glomerata* and its epiphytic microbiome dominate summer biomass in the Eel River, a Northern California river under Mediterranean (summer drought, winter rain) seasonality. Green *Cladophora* streamers proliferate in early summer, then change to yellow and then red-brown as epiphyte loads increase. We characterized successional changes in epiphytic bacteria on *Cladophora*, examining both community composition and growth rates, using quantitative Stable Isotope Probing (qSIP) and16S rRNA gene amplicon sequencing. The number of bacterial taxa increased with succession while growth rates peaked in the middle stage. NanoSIMS imaging confirmed high sulfur (S) concentrations in *Cladophora* cell walls relative to surrounding biomass, coinciding with a bloom of sulfur bacteria (bacteria that reduce or oxidize sulfur/sulfates). In general, relative abundances and growth rates were independent, indicating that either metric alone is insufficient for understanding how taxonomy and functional groups affect ecosystem processes. For instance, the relative abundance of nitrogen fixers peaked in the late summer when their relative growth rates were slowest. Such patterns may be driven by space competition limiting growth. Together, changes in abundance and relative growth rates suggest different limiting factors for different functional groups in the *Cladophora* microbiome at multiple successional stages.

## Introduction

*Cladophora glomerata* (hereafter *Cladophora)* is a green macroalga that often dominates freshwater producer biomass in temperate lakes and rivers worldwide (Zulkifly et al., 2013; Van den Hoek, 1979). It grows attached to rock or plant surfaces, with its branching filaments proliferating into streamers that can attain lengths of several meters. *Cladophora glomerata’s* rough cell walls are often covered with epiphytic diatoms, microalgae, and other microbes, and its long branching proliferations can increase stable, sunlit habitable surfaces available to these freshwater epiphytes by 4-5 orders of magnitude (Power et al., 2009, Prazukin et al., 2020, Zulkifly et al., 2012). Epiphyte attachment ultimately shades *Cladophora* and may increase drag or decrease tensile strength (Power & Marks., 2001; Graham et al., 2015; Zulkifly et al., 2013, Braus et al., 2017). While some findings suggest epiphytic organisms have a small impact on Cladophora (Dodds, 1991), little is known about the role of bacteria in the microbiome assemblage and potential function of the prokaryotic community.

In unpolluted western rivers like the Eel River in Northern California, succession among epiphytes causes *Cladophora* streamers to change color over the summer months, from green (before the host is heavily colonized), to yellow (when colonized by a mono-layer dominated by mid-successional diatoms like Cocconeis spp.), and finally to red-brown in mid-summer, a color often produced by multiple layers of the diatom *Epithemia* (Power et al., 2009; Furey et al., 2012). Diatoms are important to aquatic ecosystems since they fix C and produce polyunsaturated fatty acids (PUFAs), essential nutrients for heterotrophs (Brett et al., 2009; Dalsgaard et al., 2003). In addition, *Epithemia* species have a cyanobacterial endosymbiont that fixes nitrogen and has genes for the synthesis of 23 amino acids, many essential to animal consumers, as well as thiamine (Nakayama et al., 2014), which has been found to limit survival of salmonid fry in western North American rivers (Suffridge et al., 2024). This endosymbiont is one of several recent endosymbioses arising between N-fixing cyanobacteria and diatoms (Nakayama et al., 2011; Bandyopadhyay et al., 2011; Coale et al., 2025). In *Epithemia turgida* (Nakayama et al., 2014) and *E. clementia* (Moulin et al., 2023), the endosymbionts lack genes for both photosystems I and II, so depend entirely on host photosynthate, effectively functioning as a nitrogen-fixing organelle or “diazoplast”. *Cladophora* thus serves a key role as host to abundant *Epithemia* that serve as important food sources for heterotrophic grazers (Kupferberg, 1997, Power et al., 2009).

Although diatoms are the most visible of *Cladophora’s* epiphytes, bacteria and archaea are also important microbiome components due to their unique and significant elemental cycling contributions (Ruen-Pham et al., 2021; Graham et al., 2015; Zulkifly et al., 2012). Several bacterial phyla are ecologically important in fresh waters (*Actinobacteria, Verrucomicrobia, Bacteroidetes, Cyanobacteria*, etc.,), as they fix C and N and can decompose complex polymers (Newton et al., 2011). *Cladophora* metagenomes from the Great Lakes region have illuminated a vast array of functional genes in the microbiome (Graham et al., 2015; Byappanahalli et al., 2019; Zulkifly et al., 2013), but to date limited data exist for the bacteria and archaea living on *Cladophora* in rivers.

In freshwater systems, biological growth is often limited by N availability (Downing &McCauley, 1992; Elser et al., 2007), so microbial processes regulating N cycling strongly influence the system. N fixers, for instance, play an important role in N-limited ecosystems (Peterson & Grimm, 1992). In the Eel River, late-summer (red) stage *Cladophora* mats fix more nitrogen than previous stages (Marks et al., 2025). This might be explained by dense *Epithemia* populations, but many N-fixing bacterial taxa live in these assemblages as well (Graham et al., 2015). In addition, the bacterial community is able to perform many other functions (e.g. mercury methylation, Vitamin B12 synthesis. etc.,), highlighting the biogeochemical diversity of these systems (Tsiu et al., 2010; Graham et al., 2015; Zulkifly et al., 2012).

By measuring microbial growth and abundance at a high taxonomic resolution we can make predictions about the linkages between algal hosts and their microbiomes and element cycling processes. If organisms belonging to any of these metabolic groups are growing quickly and/or are in high abundance, then their capability to perform these functions may confer a competitive advantage resulting in high process rates (Greenlon et al., 2022). Quantitative taxon-resolved activity data can point us to processes of ecological importance at different successional stages, and which microbes drive those processes.

We used quantitative stable isotope probing (qSIP) (Hungate et al., 2015) and NanoSIMS isotopic imaging (Pett-Ridge & Weber, 2021) to identify epiphytic bacteria and archaea and measure their growth rates during the three successional stages of *Cladophora’s* microbiome in the Eel river at the Angelo Coast Range Reserve (http://angelo.berkeley.edu). We expected bacteria and archaea to increase in diversity and grow faster from each successional stage to the next, paralleling increases in eukaryotic epiphytes. We also aimed to characterize bacterial and archaeal functions, expecting more N-fixing organisms on red *Cladophora* where N-fixation is highest.

## Methods

### Quantitative Stable Isotope Probing incubations

Short *Cladophora* streamers at three different successional stages (early green, mid-successional yellow, late-successional red) were collected from the upper South Fork Eel River of northern California within the University of California, Berkeley’s Angelo Coast Range Reserve in Mendocino County (39.722468, -123.649589). The stages were defined by visual color assessment, identifying *Cocconeis* versus *Epithemia* under the microscope, and by time of season. Over the summer of 2021, green *Cladophora* and yellow *Cladophora* were collected in late May and June, red *Cladophora* was sampled in mid-July.

Samples were incubated in 98 atm% ^18^O-enriched water, with controls at natural abundance (hereafter referred to as ^16^O or control water). To replicate environmental conditions, 5 mL of river water was first collected in 15 mL falcon tubes. The tubes were dried at 60 C for 5 days until all water evaporated, leaving the previously dissolved particulate matter in the bottom of the tubes. Tufts of *Cladophora* were then collected, spun to shake off the water, and cleaned of midges and twigs with forceps. Then clumps (∼0.5 g) were allotted to each tube and resuspended along with the dried particulate matter in either 5mL of ^18^O-H_2_O or ^16^O-H_2_O. The capped sample tubes were then returned to the sites from where the algal tufts were collected and left to incubate in the running water for seven days. We incubated five ^18^O-H_2_O tubes and five ^16^O-H_2_O replicates at each algal stage for a total of 30 samples. These samples were then taken back to the lab at Northern Arizona University on dry ice for DNA extraction and qSIP analysis.

### DNA extractions

Samples were stored on dry ice and kept at -80 C before being thawed for DNA extractions. Approximately 0.2 g of algae was pulled from each sample and the water lightly shaken off. DNA was extracted using the DNeasy PowerPro Soil Kit from Qiagen, following the kit protocol. Two adjustments were made: in the first step, the silica bead tube with C1 solution and sample were bead beaten for 45 seconds twice with 5 minutes of heating at 60 C in between. In the final step the DNA on silica was heated for 2 minutes after adding the C6 solution and before centrifugation.

### Fraction separation

DNA extracts were separated into 20 to 24 density fractions using ultracentrifugation. For each sample, 5 ug of DNA, 6.85 g CsCl, gradient buffer (200 mM Tris [pH 8.0], 200 mM KCl, 2 mM EDTA), and 5 ng internal standards (adapted from Bell et al., 2023) were added to a 4.7 ml OptiSeal ultracentrifuge tube (Beckman Coulter, Inc). Tubes were then centrifuged in an Optima Max benchtop ultracentrifuge, using a Beckman TLN-100 rotor (Beckman Coulter, Inc.) at 50,000 rpm at 18°C for 72 h. Then samples were separated into 200 uL fractions and cleaned for qPCR and sequencing.

Quantitative PCR was performed in triplicate on each fraction from every sample using a BioRad CFX-384 thermal cycler. 16S rRNA genes were amplified using the EUB338F/EUB518R (Fierer et al., 2005) primer sets and 10 uL reactions containing 1 mM of Eva Green ForgetMeNot Master Mix (Thermo Fisher Scientific, Waltham MA), 0.25 mM of each primer, and 1.5 mM MgCl. Reactions were conducted using a thermocycler protocol of 95° C initial denaturation step for 2 minutes then 35 cycles of 95°C denaturation for 30 seconds, 62° C annealing for 10 seconds, and 72° C extension for 10 seconds.

### Sequencing

Sequencing was performed on an Illumina MiSeq platform in the Genetics Core Facility at Northern Arizona University. Libraries were prepared using a two-step PCR amplification of the v4 region of the 16S rRNA gene using 1 mM of each 515F and 806R sequencing primer (Parada et al, 2016), 1 mM Phusion Green master mix (Thermo Fisher Scientific, Waltham MA), and 1.5 mM MgCl_2_ for the first reaction. Then the duplicate reaction products were pooled and diluted tenfold to be used as the template DNA for the indexing PCR reaction. Here, the same 10 uL reaction was performed for each sample but using 515F and 806R indexing primers including illumina flowcell adapter sequences and a 12 nucleotide Golay barcode. Both of the PCR reactions were performed using 95 C Initial denaturation step for 2 minutes then 20 cycles (10 for the second PCR reaction) of 95 C denaturation for 30 seconds, 62 C Annealing for 10 seconds, and 72 C Extension for 10 seconds. The products from the tailing reaction were cleaned using carboxylated SeraMag Speed Beads (Sigma-Aldrich, St. Louis, MO) according to the methods in Rohland and Reich (2012). The reactions were then pooled into a single library and recleaned with the same bead protocol and quantified using Picogreen fluorescence. The library was then loaded at 11 pM onto an Illumina MiSeq according to the manufacturer protocol.

Sequences can be found on NCBI under the accession number: PRJNA1221941.

### Bioinformatics

Sequences were denoised using DADA2 in QIIME2 (Boylen et al., 2019). Sequences were assigned taxonomy using the SILVA 132 database and the QIIME2 pre-trained classifier (Robeson et al., 2020; Bokulich et al., 2018) before chimeras were filtered out. A rarefied feature table containing all sequences and a taxonomy assignment file were both imported from QIIME2 into R for qSIP analysis.

In R, we measured the ASV richness and diversity of each replicate using the Shannon-Wiener index. Then taxa were assigned individual excess atom fraction (EAF) ^18^O values based on the difference in density among amplicon sequence variants (ASV) in labeled and unlabeled samples according to the methods in (Hungate et al., 2015; Finley, 2019). EAF values were calculated using the qsip2 package in R (jeffkimbrel.github.io/qSIP2/). First, the number of 16s rRNA gene copies for each taxon was determined by multiplying the relative abundance of gene copies of each taxon (from sequencing) in each fraction of each replicate with the total number of gene copies of all taxa in the corresponding fractions and replicates. After the gene copies for each taxon were determined, taxa that were only found in one of the isotopic treatments were filtered out. With the remaining taxa the weighted average density, difference in weighted average density between labeled and unlabeled treatments, GC content, and the EAF of the DNA within the labeled treatment were calculated (Hungate et al., 2015). Relative growth rates (RGR) for each taxon were calculated from the EAF values as outlined in Koch et al., (2018) where RGR = ^18^O EAF_taxon_ /(0.98 * 0.6 * 7). 0.98 is the ^18^O enrichment of the water in the incubations, 0.6 is the percentage of oxygen atoms in DNA that are derived from water, and 7 is the number of days of incubation. These growth rates are gross calculations meaning that they do not reflect death rates.

Using PICRUST2 (Douglas et al., 2020), we estimated the 16S rRNA copy number for each ASV. We then calculated the cell count (or absolute abundance) for each ASV by multiplying its relative abundance by the total 16S rRNA copies from qPCR and then dividing the 16S rRNA abundance by the 16S rRNA copy number. We also used PICRUST2 to assign functional genes to the detected ASVs to predict the niches of active organisms. This method is certainly no replacement for metagenomics and has been shown to have low precision in environmental samples and therefore cell counts and functional groups should be viewed as best estimates and not hard observations. We also attempted to improve on this method by cross referencing the PICRUST2 results with our own functional assignments based on genus level taxonomy. This way, PICRUST2 was used as an aid in assigning functional groups rather than being the only tool. First we searched for families and genera known to have certain functions (*Azotobacter* and *Nostoc* are known N fixers for instance) (Aasfar et al., 2021; Smercina et al., 2019; Stal, 2015, Carareto Alves et al., 2014). These organisms would be assigned to the N-fixer group regardless of the PICRUST2 results. Secondly, we double checked the PICRUST2 results by cross referencing organisms that would be assigned a functional group based on PICRUST2 with genomes from the Kegg database (Kanehisa et al., 2023). Taxa that were not assigned down to a family name were not assigned to a functional group except for *Cyanobacteria*.

Statistical analysis was performed in R using one way ANOVAs and Tukey multiple comparison of means tests. When calculating means and performing statistical tests on growth rates between stages the growth rates of each bacterial population were weighted by relative abundance.

### NanoSIMS analysis

Algae tufts were preserved in 3.7% glutaraldehyde for 30 min, embedded in LR white resin following the manufacturer’s instructions (Ted Pella, # 18182), and sectioned with an ultramicrotome to produce 1 µm thick sections (Leica Ultracut). The sections were placed on glass slides, coated with approximately 10 nm of gold, and mounted in holders for NanoSIMS analysis (Pett-Ridge & Weber, 2021). Secondary ion mass spectrometry (SIMS) was performed at Lawrence Livermore National Laboratory using a Cameca NanoSIMS 50. A ∼2 pA Cs^+^ primary ion beam was focused to a ∼150 nm spot size and rastered over the sample in a 256 × 256 pixel grid to generate secondary ions for quantification. Dwell time was 1 ms/pixel, and the raster size was 40×40 microns. The secondary mass spectrometer was tuned for ∼6800 mass resolving power (1.5x correction; Pett-Ridge & Weber, 2021) to resolve isobaric interferences. ^12^C_2_ ^-, 12^C^13^C^-, 12^C^14^N^-, 12^C^15^N^-^, and ^32^S^-^ were detected in simultaneous collection mode by pulse counting on electron multipliers to generate 6 serial quantitative secondary ion images (layers). L’image data processing software (https://sites.google.com/carnegiescience.edu/limagesoftware/home) was used for data processing and visualization. Because the samples were from a separate ^13^C-labeling experiment,^32^S^-^ data were normalized to [^12^C_2_ ^-^ + ^12^C^13^C^-^/2], where the factor of two corrects for the ^12^C^13^C_2_ ^-^ count rate relative to ^12^C_2_^-^ (Pett-Ridge & Weber, 2021). Ratio images of ^32^S^-^:C were then created to detect relative S concentrations in the consortia.

## Results

### Community composition

Bacterial epiphyte assemblages differed among early (green), middle (yellow), and late (red) successional stages of the *Cladophora* microbiome. Species richness of bacteria and archaea in *Cladophora*’s microbiome significantly increased with successional stage, from green (average of 780 distinct amplicon sequence variants (ASVs)) to yellow (962 ASVs) to red (1,771 ASVs), P-value = 0.0002 (Table 1). Shannon-Wiener diversity (P-value = 2.13E-06) and evenness (P-value = 5.83E-05) measurements also increased from stage to stage, most steeply between the yellow and red stages (Figure 1, Table 1). Most (64%) ASVs detected during the red stage were unique to that stage. Stage-specific ASVs were rare on yellow (45%) and green (24%) *Cladophora* (Figure 1).

**Table 1:** Changes in community compositions of bacterial epiphytes on Cladophora in the Eel River. ASV richness, diversity, relative abundance and relative growth rates for 16s amplicons at three stages of succession. Green, yellow, and red refer to the early, middle, and late successional stages respectively. Metabolic functional groups were determined using PICRUSt2, KEGG, and confirmed in the literature. Not all ASVs belong to a functional group but all sequenced amplicons are included in the whole community richness, growth rate and diversity indices.

| Stage | Metabolism | Richness (Hmax) | standard deviation | Shannon (H) | standard deviation | Evenness (E) | Relative abundance | standard deviation | Average growth rate (new 16s copies/total 16s copies per day) | Estimated cell counts (calculated from ASV copy numbers predicted by picrust) | standard deviation |
| --- | --- | --- | --- | --- | --- | --- | --- | --- | --- | --- | --- |
| Green | Total | 780.00 | 314.56 | 4.33 | 0.26 | 0.65 | 1.00 |  | 0.07 | 1.42E+10 | 9.40E+08 |
| Yellow | Total | 961.60 | 220.67 | 4.40 | 0.42 | 0.64 | 1.00 |  | 0.09 | 2.77E+10 | 5.23E+08 |
| Red | Total | 1,770.80 | 223.97 | 5.96 | 0.18 | 0.80 | 1.00 |  | 0.07 | 1.05E+10 | 8.90E+08 |
| Green | S Reducing bacteria | 6.00 | 5.20 | 0.83 | 0.57 | 0.46 | 0.01 | 0.00 | 0.08 | 6.39E+07 | 3.34E+07 |
| Yellow | S Reducing bacteria | 4.20 | 1.10 | 0.72 | 0.31 | 0.50 | 0.00 | 0.00 | 0.13 | 2.75E+06 | 1.55E+06 |
| Red | S Reducing bacteria | 31.40 | 15.70 | 1.88 | 0.69 | 0.55 | 0.01 | 0.01 | 0.10 | 1.19E+08 | 1.26E+08 |
| Green | S Oxidizing bacteria | 49.80 | 8.47 | 2.96 | 0.12 | 0.76 | 0.08 | 0.02 | 0.08 | 9.75E+08 | 2.18E+08 |
| Yellow | S Oxidizing bacteria | 57.40 | 10.64 | 2.85 | 0.26 | 0.70 | 0.07 | 0.01 | 0.12 | 1.61E+09 | 3.27E+08 |
| Red | S Oxidizing bacteria | 80.20 | 11.97 | 3.60 | 0.18 | 0.82 | 0.09 | 0.01 | 0.09 | 1.08E+09 | 1.45E+08 |
| Green | Nitrifiers | 2.20 | 1.79 | 0.58 | 0.59 | 0.73 | 0.00 | 0.00 | 0.00 | 1.03E+07 | 1.80E+07 |
| Yellow | Nitrifiers | 3.60 | 1.67 | 0.41 | 0.24 | 0.32 | 0.00 | 0.00 | 0.06 | 4.85E+06 | 5.01E+06 |
| Red | Nitrifiers | 14.80 | 1.64 | 1.92 | 0.13 | 0.71 | 0.01 | 0.00 | 0.02 | 1.01E+08 | 1.88E+07 |
| Green | N fixers | 43.00 | 20.53 | 2.71 | 0.31 | 0.72 | 0.04 | 0.01 | 0.08 | 5.47E+08 | 1.08E+08 |
| Yellow | N fixers | 52.20 | 10.69 | 2.87 | 0.44 | 0.72 | 0.03 | 0.01 | 0.09 | 6.57E+08 | 1.65E+08 |
| Red | N fixers | 92.00 | 22.11 | 3.17 | 0.42 | 0.70 | 0.12 | 0.03 | 0.05 | 9.27E+08 | 1.56E+08 |
| Green | Fe Reducing bacteria | 6.00 | 2.92 | 1.34 | 0.15 | 0.75 | 0.01 | 0.00 | 0.09 | 1.14E+08 | 6.16E+07 |
| Yellow | Fe Reducing bacteria | 7.20 | 1.30 | 0.66 | 0.26 | 0.33 | 0.03 | 0.01 | 0.13 | 2.78E+08 | 7.36E+07 |
| Red | Fe Reducing bacteria | 13.40 | 6.23 | 1.56 | 0.52 | 0.60 | 0.01 | 0.01 | 0.12 | 1.66E+08 | 6.53E+07 |
| Green | Fe Oxidizing bacteria | 8.40 | 2.41 | 1.34 | 0.17 | 0.63 | 0.01 | 0.00 | 0.07 | 1.14E+08 | 6.16E+07 |
| Yellow | Fe Oxidizing bacteria | 10.00 | 1.87 | 1.30 | 0.31 | 0.56 | 0.01 | 0.00 | 0.09 | 2.78E+08 | 7.36E+07 |
| Red | Fe Oxidizing bacteria | 13.00 | 2.35 | 2.07 | 0.21 | 0.81 | 0.02 | 0.00 | 0.08 | 1.66E+08 | 6.53E+07 |
| Green | Denitrifiers | 14.20 | 2.39 | 1.68 | 0.13 | 0.63 | 0.02 | 0.00 | 0.10 | 6.36E+08 | 2.13E+08 |
| Yellow | Denitrifiers | 14.40 | 1.14 | 1.78 | 0.29 | 0.67 | 0.03 | 0.02 | 0.13 | 9.03E+08 | 5.36E+08 |
| Red | Denitrifiers | 14.60 | 5.13 | 2.03 | 0.51 | 0.76 | 0.02 | 0.01 | 0.12 | 1.52E+08 | 4.64E+07 |
| Green | Cyanobacteria | 32.60 | 19.37 | 2.45 | 0.33 | 0.70 | 0.02 | 0.03 | 0.05 | 1.27E+08 | 2.40E+08 |
| Yellow | Cyanobacteria | 48.60 | 17.44 | 2.38 | 0.32 | 0.61 | 0.01 | 0.00 | 0.03 | 1.31E+08 | 7.81E+07 |
| Red | Cyanobacteria | 91.20 | 15.56 | 3.16 | 0.07 | 0.70 | 0.15 | 0.05 | 0.03 | 9.02E+08 | 2.55E+08 |
| Green | Decomposing phyla | 80.40 | 25.30 | 3.18 | 0.34 | 0.72 | 0.02 | 0.02 | 0.03 | 2.15E+08 | 2.30E+08 |
| Yellow | Decomposing phyla | 153.40 | 40.25 | 3.57 | 0.39 | 0.71 | 0.03 | 0.01 | 0.05 | 8.43E+08 | 4.29E+08 |
| Red | Decomposing phyla | 354.80 | 39.81 | 4.80 | 0.15 | 0.82 | 0.14 | 0.03 | 0.06 | 1.50E+09 | 3.62E+08 |

**Figure 1.**
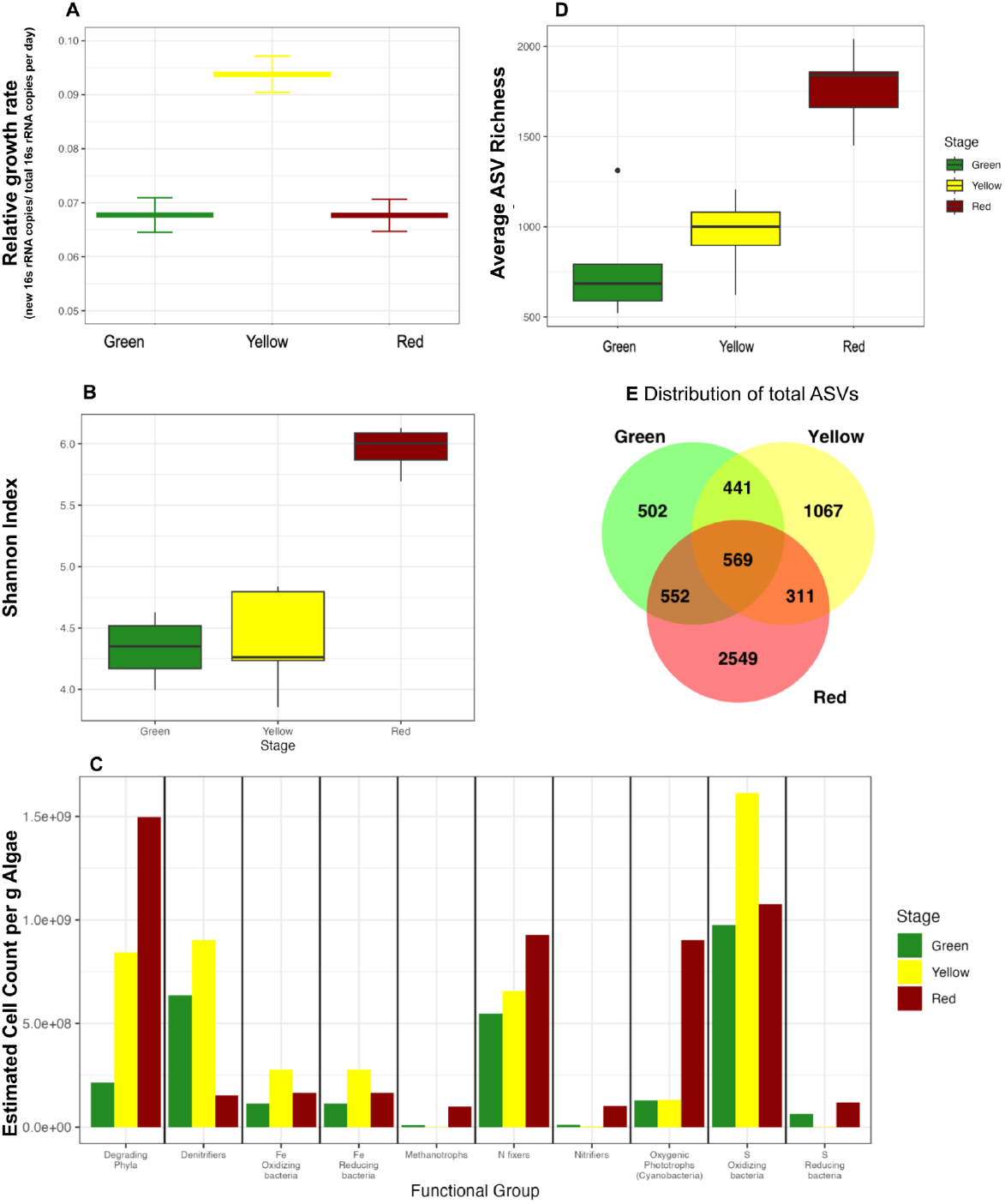
Community changes in abundance and growth rates between Cladophora epiphyte successional stages. **A)** The average relative growth rate of the assemblage at each stage (P-value <2e-16). Green is early summer, yellow is mid summer and red is late summer. Bars represent the average growth rate of all populations sampled at each site and error bars represent the 95% confidence interval calculated from the mean and variance of taxon specific growth rates weighted by relative abundance. **B)** The Shannon-Weiner diversity index across stages (P-value = 2.13e-06). **C)** Absolute abundances (Calculated using PiCRUSt) of N fixers (P-value = 8.12e-06), S reducers (0.07), decomposers (4.3e-06), Cyanobacteria (2.53e-05), and nitrifiers (9.94e-08) increased on red Cladophora while the other functional groups had the highest abundances on yellow (P-values = 0.0163, 0.0106, 0.0113, 0.393 for denitrifiers, Fe reducers, Fe oxidizers and S oxidizers respectively). **D).** ASV richness (P-value = 0.000118). **E)** Venn Diagram of all detected ASVs across the successional stages.

### Growth rates

On average, bacterial populations exhibited faster growth on yellow *Cladophora* (0.094 new 16S rRNA gene copies/total 16S rRNA gene copies per day), compared to 0.068 d^-1^ for both red and green stages (Figure 1A, Table 1). A Tukey multiple comparisons of means confirmed that the average yellow stage growth rate was significantly different than the green and red stages (Yellow - Green: P-value < 2E-16; Yellow-Red: P-value < 2E-16). Phylum was a strong predictor of growth rate (Figure 2B, C, and D)—fast growing taxa on one stage had similarly rapid relative growth on other *Cladophora* stages. When growth rates of individual populations differed among stages, they tended to be slower on green (P-value < 2E-16) than on red or yellow *Cladophora* (Figure 2). The fastest growing taxa occurred predominantly on the yellow and red stage *Cladophora* (Figure 1A). All functional groups peaked in growth rate on yellow *Cladophora* except for the oxygenic phototrophs (Cyanobacteria). The average growth rate of photosynthetic bacteria was fastest on green *Cladophora* and slowed at each subsequent successional stage but not significantly.

**Figure 2:**
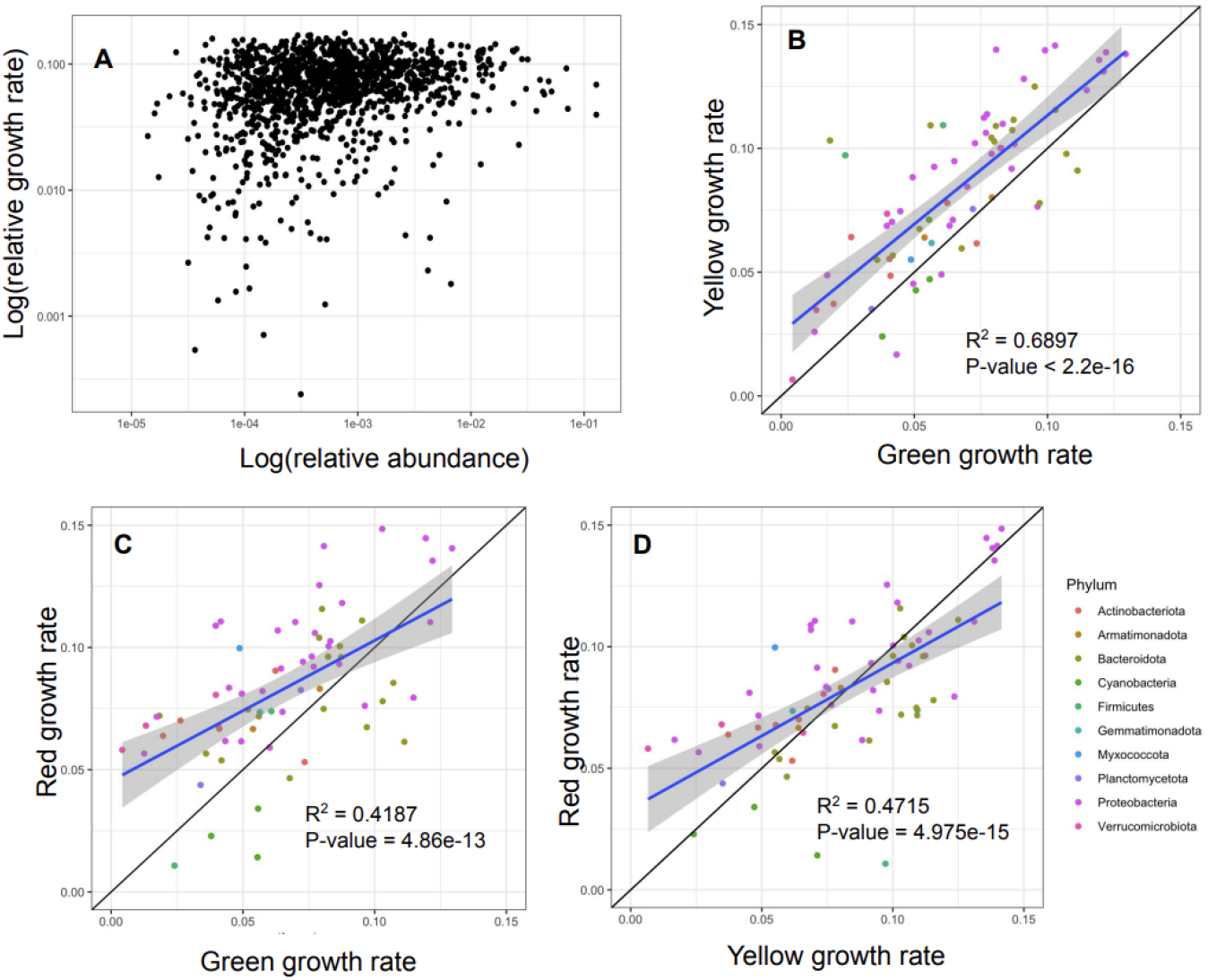
Comparison of taxon specific bacterial growth rates from the Cladophora microbiome in the Eel River (Mendocino CA). A) Each point is the relative growth rate (y axis) and relative abundance (x axis) of a taxon on a given successional stage of *Cladophora*. Growth rates and abundances are poorly correlated (R^2^ = 0.00318, P-value = 0.01565), a negative correlation is expected if cell growth slows due to intraspecific competition as they reach higher densities. B-D) Populations of bacteria that occur across three successional stages (green, yellow, and red for early, middle and late stages) tend to grow at similar rates across stages.

### Decomposing Bacteria

Heterotrophic bacteria use organic material for their carbon source. There are a few phyla that are particularly adept at breaking down polymers. The phyla *Verrucomicrobiota, Plantmycetota*, and *Actinobacteria* produce enzymes that break down algal cell walls. These decomposers were the only groups that increased in both abundance (with red relative abundances being significantly higher than the other two stages: P-value = 4.3E-06) and relative growth rate (P-value = 3.374E-07) through time. On the red *Cladophora*, these phyla made up almost 14% of the prokaryotic 16S rRNA gene copies. They increased from 2.15E+08 cells per gram algae on green *Cladophora* to 8.43E+08 cells on yellow to 1.50E+09 cells on red. In addition to this increase in abundance, the average relative growth rate of the whole community almost doubled from 0.03 new 16s rRNA copies/ total 16s rRNA copies per day on green *Cladophora* to 0.06 new 16s rRNA copies/ total 16s rRNA copies per day on red *Cladophora*.

### Nitrogen Fixers

The abundance of N-fixers increased 4-fold between the yellow and red stages, from 3% to 13% of the community abundance (P-value = 8.12E-06) see (Table 1). By the red stage, N-fixers were the most abundant and most species rich group, surpassing the sulfate oxidizing bacteria and second only to oxygenic phototrophs (all free-living *Cyanobacteria)*, many of which are N fixers. In fact, while free living Cyanobacteria were more abundant than N-fixers on red Cladophora (late stage), the N-fixers had faster growth rates during this season (S1, P-value = 0.03).

Among the N-fixers on the green *Cladophora*, only *Proteobacteria, Firmicutes, Cyanobacteria* and one ASV belonging to *Campilobacterota* had positive growth rates. Of these, *Proteobacteria* grew the fastest, with the fastest-growing ASV classified in the *Rhodocyclaceae* family. The composition and growth of the N-fixing assemblage changed from stage to stage. On red *Cladophora*, growing *Cyanobacteria* became far more abundant, with free-living and endosymbiotic taxa together making up 45% of the N-fixers with detectable growth, compared to 8% on green (Table 1). The proteobacteria were still the fastest growing group however, with the fastest growing *Cyanobacteria* (*Nostocaceae*) growing at less than a third of the rate of the fastest growing *Proteobacteria* (Figure S2). N fixers also became more diverse. The Shannon Wiener index increased from 2.71 on green to 3.17 on red (Table 1), with new phyla (*Desulfobacteria*, one Archaeal taxon, and one *Myxococcota*) growing on the red *Cladophora*.

The most abundant nitrogen fixing ASV on red *Cladophora* (accounting for 25% of all N fixers) was an N-fixing endosymbiont descended from *Cyanothece* (Nakayama et al., 2011). This endosymbiont (or diazoplast (Moulin et al., 2024)) is found in the diatom genus *Epithemia*, which is very abundant on red *Cladophora*.

### Fe, S, N Reducing Bacteria

Bacteria capable of dissimilatory reduction of N, Fe, and S were present and growing across all stages of algal development. Denitrifying bacteria were the most abundant at each stage. Iron reducers peaked in abundance on yellow *Cladophora* and sulfate reducers peaked on red *Cladophora*. Denitrifying bacteria decreased in abundance from stage to stage but peaked in growth rate on yellow *Cladophora*. Sulfur and Fe reducers also grew fastest at that stage, mirroring the pattern of the whole community however only Fe reducers had growth rates that were statistically correlated to successional stage (P-value = 0.0042).

### Fe, S, N Oxidizing Bacteria

Organisms that are capable of oxidizing reactions such as nitrification, iron oxidation and sulfur oxidation to sulfate also increased in growth and relative abundance between the green and red algal stages. The most abundant oxidizers here were the sulfur oxidizers which were also the most abundant group in the green and yellow stages. The other oxidizing bacteria were rare during the first two stages with nitrifiers representing less than 0.1% of the community. Like the rest of the community, the oxidizing bacteria tended to grow slowest in the green stage and fastest in the yellow stage (Table 1, S1, S2). Nitrifiers and iron oxidizers did not have statistically significant changes in growth between successional stages but sulfur oxidizers increased significantly in growth rate between the green and yellow stages, and decreased significantly from yellow to red (P-value = 5.24E-06, S1, S2).

### Sulfur and Cladophora

During the late/red successional stage we detected relatively fast growth of both S reducers and S oxidizers compared to other groups (Table 1, S1) with significant differences in growth rates between these sulfur bacteria and *Cyanobacteria*, degrading phyla, and nitrifiers (P-values: <2E-16, <2E-16, and 0.001 respectively for oxidizers and 0.0007, 0.0224, and 0.01 respectively for reducers). Coinciding with the peak of these bacteria during the late successional stage is the senescence of *Cladophora* which has a cell wall found to be enriched in S relative to the surrounding community (Figure 3). The NanoSIMS image, taken from a sample of red (late stage) *Cladophora* shows high sulfur concentrations in the cell wall relative to the cytoplasm as well and that this sulfur is not uniformly distributed in this cross section.

**Figure 3:**
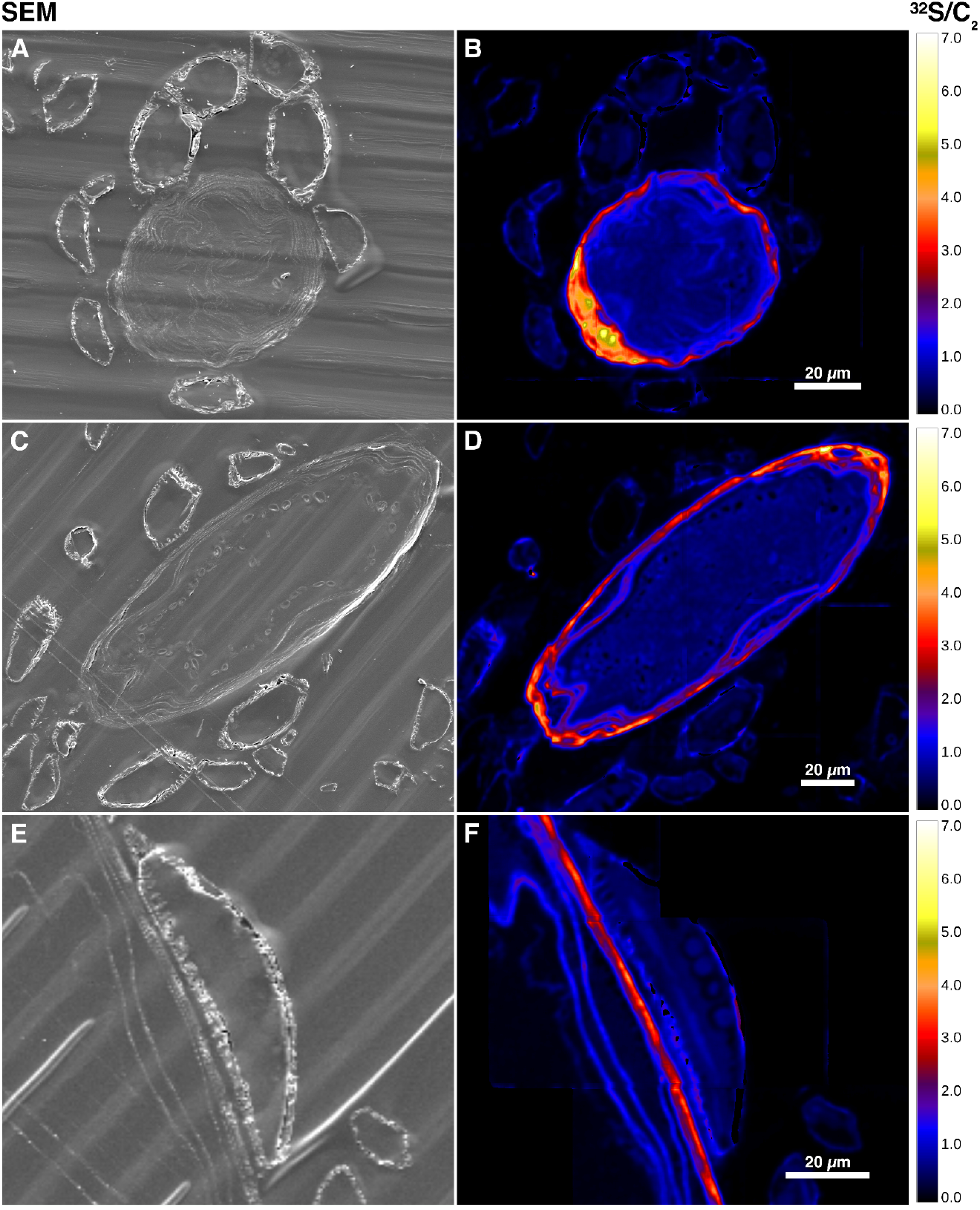
A cross-section of three Cladophora filaments and surrounding epiphytes. **A)** Scanning Electron Microscope image of a *Cladophora* cell (largest circular structure). The surrounding cells are likely *Epithemia*. **B)** NanoSIMS image of the same section. ^32^S counts are normalized to total C meaning that high ^32^S/C (shown by more yellow coloring) indicates high S content in that area relative to total cell C content.

## Discussion

A population’s impact on an ecosystem depends on both abundance and per capita rates at which individuals perform impactful activities. The latter are likely to correlate with the population’s growth rate. Population impacts can also vary with functional characteristics of the species and strain, across environmental contexts or gradients. We describe the potential impact of several functional groups in the *Cladophora* microbiome based on their abundances and relative growth rates. Most functional groups increased in abundance over succession and peaked in relative growth rate on yellow *Cladophora*. By the red stage, the microbiome was dominated by decomposers, N fixers, *Cyanobacteria*, and sulfur bacteria.

Bacteria with N-fixing capabilities had low growth rates (especially on red *Cladophora*) but increased in ASV richness and absolute and relative abundance. Nitrogen fixers were the most abundant functional group on red *Cladophora* largely due to an increase in the abundance of nitrogen fixing *Cyanobacteria* and *Epithemia’s* diazoplast. The diazoplast cannot be used to predict diatom growth rates or abundances due to variation in diazoplast numbers among different host diatom cells (Hagino et al., 2013., Marks et al., 2025), their increase indicates that fixation is an important source of nitrogen during late succession, as rates and *Epithemia* diatoms both increase in the mainstem river (DeYoe et al., 1992., Power et al., 2009). Since N-fixation is an energetically costly process, the slow growth observed on red *Cladophora* may be a result of these organisms allocating resources to fix nitrogen (Dutkiewicz et al., 2012). Many nitrogen fixers (especially *Cyanobacteria*) are facultative, meaning that they only fix nitrogen when they are starved for the element (Herrero et al, 2001; Pearl, 2017). The decline in the growth rates of this group on red *Cladophora*, while their abundances remained high, corresponds with a peak in N-fixation rates at the site (Marks et al., 2025). This suggests that nitrogen availability to the ecosystem dwindled between early and mid-summer.

In previous studies during the summer drought, watershed inputs of nitrogen to our study site along the upper South Fork Eel River were minimal and concentrations of dissolved inorganic N were very low, yet concentrations of dissolved organic N increased (Finlay et al., 2011). Finlay et al. (2011, see also Power et al., 2009) deduced that DON increased in later summer due to riverine N-fixation. Inorganic NH_4_ is lost to uptake by biota or to nitrification (oxidation to NO_3_) and then to gaseous N_2_O or N_2_ (denitrification) (Xia et al., 2018). We found few nitrifiers present and growing on the *Cladophora*, but denitrifying bacteria were one of the fastest growing functional groups and were the most abundant in early season *Cladophora* (Table 1). Denitrification is not always an obligate function for bacteria, and growth does not directly indicate denitrification, but the fast growth of organisms with this capability could mean that denitrification on green *Cladophora* could be a factor driving nitrogen limitation during later successional stages, and setting up conditions that favor nitrogen fixers.

Fe and S cycling organisms grew rapidly – growth rates of both oxidizers and reducers increased from green and red *Cladophora* and peaked on yellow. This suggests that reducible substrates must have been readily available, and that hypoxic or anaerobic microsites were present within the *Cladophora* branches. Fe is needed for chloroplasts and nitrogenase enzymes, so, like N, often limits primary productivity in streams (Lis et al., 2015; Lau et al., 2016). The growth of organisms that can reduce Fe may facilitate increases in the *Cyanobacteria* population given that Fe limitation can reduce the growth of phytoplankton (Larson et al., 2018). Given that late summer N availability in the Eel may be driven by N-fixation, the processing of S and Fe could be important factors in N cycling. Fe is often derived from sediments and upwelling of bioavailable Fe has been linked to cyanobacterial blooms (Dengg et al., 2023). Sulfur is also an essential element for microbial enzymes and amino acids. The byproducts and intermediates of sulfur redox reactions (both reduction and oxidation) are biogeochemically important (Vigneron et al., 2021). Sulfate reduction by bacteria can recycle S in sediments (Bowles et al., 2014). Both S oxidizing and S reducing bacteria had their highest relative abundances on red *Cladophora*, where together they made up over 10% of the prokaryotic community.

The increase of S cycling organisms during the red stage could be due to the decay of *Cladophora. Cladophora’s* cell wall is enriched in sulfated polysaccharides (Figure 3) which have been shown to protect *Cladophora* from osmotic stress and superoxide (Arata et al., 2017) and may have bactericidal, antifungal, and antiviral properties (Jasem et al., 2023). *Planctomycetota* and *Verrucomicrobiota* (two phyla that increased in abundance over the summer) produce sulfatases which they use to degrade sulphated polysaccharides (Orellana et al, 2022; Klimek et al, 2024). Sulfur reducing bacteria grew particularly quickly on red senescing *Cladophora*, (Table 1, Figure 1), and may have been responsible for the lack of uniformity in the sulfur content of the exposed *Cladophora* wall in figure 3. Other studies have documented purple sulfur bacteria on the surface of *Cladophora* (Olapade et al., 2006; Wang et al., 2023) and these sulfur degrading bacteria likely play a crucial role in the recycling of sulfate derived from *Cladophora* cell walls.

In addition to the sulfur cyclers, red *Cladophora* was also rich in organisms that promote decomposition in streams. Decomposers were the only group to increase in both abundance and growth rate between yellow and red stages. Organisms that are known to decompose complex algal and plant polymers are common epiphytes on algal hosts (Seyedsayamdost et al, 2011; Abate et al., 2024). Many taxa within these groups are beneficial to algal hosts by competing for resources with pathogens, producing essential vitamins, and exchanging nutrients (Cirri & Pohnert, 2019). We found many organisms growing at all stages of the *Cladophora* that are likely involved in decomposition processes after algal death. That they are growing and present early is an indication that these organisms may perform early symbiotic functions. *Roseobacter* (detected on the *Cladophora* mats as well) has been shown to be both a mutualistic epiphyte and a decomposer on other eukaryotic algal types (Seyedsayamdost et al., 2011; Segev et al., 2016). On *Cladophora* bacteria may donate vitamins and nutrients to the host algae (Graham et al., 2015), but while there is good understanding of bacterial communities and functional groups that occur on *Cladophora*, measuring fluxes and algal exudates in these consortia would be the next step in deciphering algal-bacterial interactions in this system.

## Conclusions

To our knowledge, this is the first study to measure taxon-specific growth rates in situ during bacterial succession in an algal microbiome. A taxon’s abundance and population growth rate, while not the only factors that make a population important, mediate its impact on an ecosystem and are often not correlated. For instance, *Cyanobacteria* and N-fixing bacteria grew more slowly from stage to stage despite becoming more abundant (possibly reflecting their population’s nearing its carrying capacity). Organisms that were present across all stages tended to have consistent growth rates from stage to stage, however most taxa grew faster late in the summer than the early successional stages (Figure 2). Although N-fixers had lower than average growth rates, they were abundant on the red *Cladophora* where N-fixation rates have been measured to be the highest (Marks et al., 2025). Conversely, Fe, S and N reducing bacteria were relatively sparse, but consistently had growth rates that were faster than the community average. Low abundance organisms may therefore have disproportionate per capita effects on biochemical processes in the ecosystem.

Bacteria capable of varying biochemical processes also changed in growth and abundance over succession. Decomposers, N-fixers, *Cyanobacteria*, and S cyclers all were dominant functions on the red stage when on earlier stages all functional groups were more similar to each other in growth and abundance. In this system succession happens during a warming and drying summer meaning that the early stage algae sit at cooler temperatures and experience a higher water flow. As the summer proceeds the waterline lowers and the air and water temperatures increase. With warmer and drier summers in northern california we may expect that these successional changes will happen faster (Power et al., 2024). Further work on how the microbiome responds to these environmental changes would contribute both to our understanding of aquatic ecology but also in predicting and preventing algal blooms both in the Eel river and in other waterways.

## Acknowledgements

We thank Sean Furey and Christina Ramon for help in the lab and Philip Georgakakos and Peter Steel for insights into the site and the University of California Natural Reserve System for providing a protected site and infrastructural support for this research. This work was funded by the National Science Foundation grant no (2125088), with other support from the National Science Foundation CZP EAR-1331940 for the Eel River Critical Zone Observatory. Work at LLNL was conducted under the auspices of the US Department of Energy under Contract DE-AC52-07NA27344.

## References

1. Zulkifly, S., Graham, J. M., Young, E. B., Mayer, R. J., Piotrowski, M. J., Smith, I., & Graham, L. E. (2013). The Genus Cladophora Kützing (Ulvophyceae) as a Globally Distributed Ecological Engineer. Journal of Phycology, 49(1), 1–17.

2. Van den Hoek C. (1979). The phytogeography of Cladophora (Chlorophyceae) in the Northern Atlantic Ocean, in comparison to that of other benthic algal species. Helgoländer Wiss. Meeresunters, 32, 374–393. doi: 10.1007/BF02189592.

3. Prazukin, A., Shadrin, N., Balycheva, D., Firsov, Y., Lee, R., & Anufriieva, E. (2021). Cladophora spp. (Chlorophyta) modulates the environment and creates a habitat for microalgae in hypersaline waters. Europeak Journal of Phycology, 56(3), 231–243. doi:10.1080/09670262.2020.1814423

4. Zulkifly S., Hanshew A., Young E.B., Lee P., Graham M.E., Graham M.E., Piotrowski M., Graham L.E. (2012). The epiphytic microbiota of the globally widespread macroalga Cladophora glomerata (Chlorophyta, Cladophorales). Am J Bot, 99(9), 1541–52. doi: 10.3732/ajb.1200161.

5. Power, M.E., Lowe, R., Furey, P.C., Welter, J., Limm, M., Finlay, J., Bode, C., Chang, S., Goodrich, M., Sculley, J. (2009). Algal mats and insect emergence in rivers under Mediterranean climates: Towards photogrammetric surveillance. Freshwater Biology, 54, 2101–2115.

6. Marks, J. C., & Power, M. E. (2001). Nutrient induced changes in the species composition of epiphytes on Cladophora glomerata Kütz. Hydrobiologia, 450, 187–196. 10.1023/A:1017596927664

7. Graham, L. E., Knack, J. J., Graham, M. E., Graham, J. M., & Zulkifly, S. (2015). A metagenome for lacustrine Cladophora (Cladophorales) reveals remarkable diversity of eukaryotic epibionts and genes relevant to materials cycling. Journal of Phycology, 51(3), 408–418. doi:10.1111/jpy.12296

8. Braus, M. J., Graham, L. E., & Whitman, T. L. (2017). Spatiotemporal dynamics of the bacterial microbiota on lacustrine Cladophora glomerata (Chlorophyta). Journal of Phycology, 53(6), 1255–1262. doi:10.1111/jpy.12573

9. Dodds WK. (1991). Community interactions between the filamentous alga Cladophora glomerata (L.) Kuetzing, its epiphytes, and epiphyte grazers. Oecologia, 85(4), 572–580. doi: 10.1007/BF00323770. PMID: 28312505.

10. Furey, P. C., Lowe, R. L., Power, M. E., & Campbell-Craven, A. M. (2012). tMidges, Cladophora, and epiphytes: shifting interactions through succession. Society for Freshwater Science, 31(1), 93–107.

11. Brett, M. T., Kainz, M., Taipale, S. J., Seshan, H. (2009). tPhytoplankton, not allochthonous carbon, sustains herbivorous zooplankton production. Proceedings of the National Academy of Sciences USA, 106, 21197–21201.

12. Dalsgaard J., John, M.S., Kattner G., Müller-Navarra, D., Hagen, W. (2003). Fatty acid trophic markers in the pelagic marine environment. Adv. Mar. Biol, 46, 225–340. doi: 10.1016/s0065-2881(03)46005-7.

13. Nakayama, T., Ikegami, Y., Ishida, K., Inagaki, Y. & Inouye, I. (2011). Spheroid bodies in rhopalodiacean diatoms were derived from a single endosymbiotic cyanobacterium. J. Plant Res. 124, 93–97.

14. Suffridge, C.P., Shannon, K.C., Matthews, H., Johnson, R.C., Jeffres, C., Mantua, N., Ward, A.E., Holmes, E., Kindopp, J., Aidoo, M., Colwell, F.S. (2024). Connecting thiamine availability to the microbial community composition in Chinook salmon spawning habitats of the Sacramento River basin. Appl Environ Microbiol, 90:e01760–23. 10.1128/aem.01760-23

15. Bandyopadhyay, A., Elvitigala, T., Welsh, E., Stöckel, J., Liberton, M., Min, H., Sherman, L. A., & Pakrasi, H. B. (2011). Novel metabolic attributes of the genus cyanothece, comprising a group of unicellular nitrogen-fixing Cyanothece. mBio, 2(5), e00214–11. 10.1128/mBio.00214-11

16. Coale, T. H., Loconte, V., Turk-Kubo, K. A., Vanslembrouck, B., Mak, W. K. E., Cheung, S., Ekman, A., Chen, J. H., Hagino, K., Takano, Y., Nishimura, T., Adachi, M., Le Gros, M., Larabell, C., & Zehr, J. P. (2024). Nitrogen-fixing organelle in a marine alga. Science (New York, N.Y.), 384(6692), 217–222. 10.1126/science.adk1075

17. Moulin, S. L. Y., Frail, S., Braukmann, T., Doenier, J., Steele-Ogus, M., Marks, J. C., Mills, M. M., & Yeh, E. (2024). The endosymbiont of Epithemia clementina is specialized for nitrogen fixation within a photosynthetic eukaryote. ISME communications, 4(1), ycae055. 10.1093/ismeco/ycae055

18. Kupferberg, S.J. (1997), BULLFROG (RANA CATESBEIANA) INVASION OF A CALIFORNIA RIVER: THE ROLE OF LARVAL COMPETITION. Ecology, 78: 1736–1751. 10.1890/0012-9658

19. Ruen-Pham, K., Graham, L.E., Satjarak, A. (2021). Spatial Variation of Cladophora Epiphytes in the Nan River, Thailand. Plants (Basel), 10(11), 2266. doi: 10.3390/plants10112266. PMID: 34834629; PMCID: PMC8622721.

20. Newton, R. J., Jones, S. E., Eiler, A., McMahon, K. D., & Bertilsson, S. (2011). A guide to the natural history of freshwater lake bacteria. Microbiology and Molecular Biology Reviews, 75(1), 14–49.

21. Byappanahalli, M. N., Nevers, M. B., Przybyla-Kelly, K., Ishii, S., King, T. L., & Aunins, A. W. (2019). Great Lakes Cladophora harbors phylogenetically diverse nitrogen-fixing microorganisms. Environmental DNA, 1(2), 186–195. doi:10.1002/edn3.20

22. Downing, John A., McCauley, Edward. (1992), The nitrogen : phosphorus relationship in lakes, Limnology and Oceanography, 37, doi: 10.4319/lo.1992.37.5.0936.

23. Elser, J.J., Bracken, M.E.S., Cleland, E.E., Gruner, D.S., Harpole, W.S., Hillebrand, H., Ngai, J.T., Seabloom, E.W., Shurin, J.B. and Smith, J.E. (2007), Global analysis of nitrogen and phosphorus limitation of primary producers in freshwater, marine and terrestrial ecosystems. Ecology Letters, 10: 1135–1142. 10.1111/j.1461-0248.2007.01113.x

24. Peterson, C. G., & Grimm, N. B. (1992). Temporal Variation in Enrichment Effects during Periphyton Succession in a Nitrogen-Limited Desert Stream Ecosystem. Journal of the North American Benthological Society, 11(1), 20–36. 10.2307/1467879

25. Marks, J.C., Zampini, M., Fitzpatrick, R.M., Hungate, B.A., Kariunga, S.H., Leshyk, V., Samo, T., Thomas, S., Weber, P., Wulf, M., Schwartz, E., Pett-Ridge, J., Power, M.E. (2025). Ecosystem consequences of a nitrogen fixing proto-organelle. PNAS, in revision.

26. Martin Tsz Ki Tsui, Jacques C. Finlay, Steven J. Balogh, and Yabing H. Nollet. (2010). In Situ Production of Methylmercury within a Stream Channel in Northern California. Environmental Science & Technology, 44 (18), 6998–7004. doi: 10.1021/es101374y

27. Greenlon, A., Sieradzki, E., Zablocki, O., Koch, B.J., Foley, M.M., Kimbrel, J.A., Hungate, B.A., Blazewicz, S.J., Nuccio, E.E., Sun, C.L., Chew, A., Mancilla, C.J., Sullivan, M.B., Firestone, M., Pett-Ridge, J., Banfield, J.F. (2022). Quantitative Stable-Isotope Probing (qSIP) with Metagenomics Links Microbial Physiology and Activity to Soil Moisture in Mediterranean-Climate Grassland Ecosystems. mSystems, 7(6) doi: 10.1128/msystems.00417-22.

28. Hungate, B.A., Mau, R.L., Schwartz, E., Caporaso, J.G., Dijkstra, P., van Gestel, N., Koch, B.J., Liu, C.M., McHugh, T.A., Marks, J.C., Morrissey, E.M., Price, L.B. (2015). Quantitative microbial ecology through stable isotope probing. Appl Environ Microbiol, 81(21), 7570–81. doi: 10.1128/AEM.02280-15. Epub 2015 Aug 21. PMID: 26296731; PMCID: PMC4592864.

29. Pett-Ridge J., Weber, P.K. (2022). NanoSIP: NanoSIMS Applications for Microbial Biology. Methods Mol Biol. 2349, 91–136. doi: 10.1007/978-1-0716-1585-0_6. PMID: 34718993.

30. Fierer N., Jackson, J.A., Vilgalys, R., Jackson, R.B. (2005). Assessment of soil microbial community structure by use of taxon-specific quantitative PCR assays. Appl Environ Microbiol. 71(7), 4117–20. doi: 10.1128/AEM.71.7.4117-4120.2005. PMID: 16000830; PMCID: PMC1169028.

31. Parada, A. E., Needham, D. M., & Fuhrman, J. A. (2016). Every base matters: assessing small subunit rRNA primers for marine microbiomes with mock communities, time series and global field samples. Environmental Microbiology, 18(5), 1403–1414

32. Rohland N, Reich D. (2012). Cost-effective, high-throughput DNA sequencing libraries for multiplexed target capture. Genome Res. 22(5), 939–46. doi: 10.1101/gr.128124.111. Epub 2012 Jan 20. PMID: 22267522; PMCID: PMC3337438.

33. Bolyen E, Rideout JR, Dillon MR, Bokulich NA, Abnet CC, Al-Ghalith GA, Alexander H, Alm EJ, Arumugam M, Asnicar F, Bai Y, Bisanz JE, Bittinger K, Brejnrod A, Brislawn CJ, Brown CT, Callahan BJ, Caraballo-Rodrïguez AM, Chase J, Cope EK, Da Silva R, Diener C, Dorrestein PC, Douglas GM, Durall DM, Duvallet C, Edwardson CF, Ernst M, Estaki M, Fouquier J, Gauglitz JM, Gibbons SM, Gibson DL, Gonzalez A, Gorlick K, Guo J, Hillmann B, Holmes S, Holste H, Huttenhower C, Huttley GA, Janssen S, Jarmusch AK, Jiang L, Kaehler BD, Kang KB, Keefe CR, Keim P, Kelley ST, Knights D, Koester I, Kosciolek T, Kreps J, Langille MGI, Lee J, Ley R, Liu YX, Loftfield E, Lozupone C, Maher M, Marotz C, Martin BD, McDonald D, McIver LJ, Melnik AV, Metcalf JL, Morgan SC, Morton JT, Naimey AT, Navas-Molina JA, Nothias LF, Orchanian SB, Pearson T, Peoples SL, Petras D, Preuss ML, Pruesse E, Rasmussen LB, Rivers A, Robeson MS 2nd, Rosenthal P, Segata N, Shaffer M, Shiffer A, Sinha R, Song SJ, Spear JR, Swafford AD, Thompson LR, Torres PJ, Trinh P, Tripathi A, Turnbaugh PJ, Ul-Hasan S, van der Hooft JJJ, Vargas F, Vázquez-Baeza Y, Vogtmann E, von Hippel M, Walters W, Wan Y, Wang M, Warren J, Weber KC, Williamson CHD, Willis AD, Xu ZZ, Zaneveld JR, Zhang Y, Zhu Q, Knight R, Caporaso JG. (2019). Reproducible, interactive, scalable and extensible microbiome data science using QIIME 2. Nat Biotechnol. 37(8):852–857. doi: 10.1038/s41587-019-0209-9. Erratum in: Nat Biotechnol. 2019 Sep;37(9):1091. PMID: 31341288; PMCID: PMC7015180.

34. Robeson II, M.S., O’Rourke, D.R., Kaehler, B.D., Ziemski, M., Dillon, M.R., Foster, J.T., Bokulich, N.A. RESCRIPt: Reproducible sequence taxonomy reference database management for the masses. bioRxiv 2020.10.05.326504; doi: 10.1101/2020.10.05.326504

35. Bokulich, N. A., Kaehler, B. D., Rideout, J. R., Dillon, M., Bolyen, E., Knight, R., Huttley, G. A., & Gregory Caporaso, J. (2018). Optimizing taxonomic classification of marker-gene amplicon sequences with QIIME 2’s q2-feature-classifier plugin. Microbiome, 6(1), 90. 10.1186/s40168-018-0470-z

36. Finley, B. K., Hayer, M., Mau, R. L., Purcell, A. M., Koch, B. J., van Gestel, N. C., Schwartz, E., & Hungate, B. A. (2019). Microbial Taxon-Specific Isotope Incorporation with DNA Quantitative Stable Isotope Probing. Methods in molecular biology (Clifton, N.J.), 2046, 137–149. 10.1007/978-1-4939-9721-3_11

37. Douglas, G. M., Maffei, V. J., Zaneveld, J. R., Yurgel, S. N., Brown, J. R., Taylor, C. M., Huttenhower, C., & Langille, M. G. I. (2020). PICRUSt2 for prediction of metagenome functions. Nature biotechnology, 38(6), 685–688. 10.1038/s41587-020-0548-6

38. Aasfar, A., Bargaz, A., Yaakoubi, K., Hilali, A., Bennis, I., Zeroual, Y., & Issam Meftah Kadmiri. (2021). Nitrogen Fixing Azotobacter Species as Potential Soil Biological Enhancers for Crop Nutrition and Yield Stability. Frontiers in Microbiology, 12. 10.3389/fmicb.2021.628379

39. Smercina, D.N., Evans, S.E., Friesen, M.L., Tiemann, L.K. (2019). To Fix or Not To Fix: Controls on Free-Living Nitrogen Fixation in the Rhizosphere. Applied Environmental Microbiology, 85(6). 10.1128/AEM.02546-18

40. Stal, Lucas J. (2015). Nitrogen Fixation in Cyanobacteria. Encyclopedia of Life Sciences. 10.1002/9780470015902.a0021159.pub2

41. Carareto Alves, L.M., de Souza, J.A.M., Varani, A.d.M., Lemos, E.G.d.M. (2014). The Family Rhizobiaceae. In: Rosenberg, E., DeLong, E.F., Lory, S., Stackebrandt, E., Thompson, F. (eds) The Prokaryotes. Springer, Berlin, Heidelberg. 10.1007/978-3-642-30197-1_297

42. Kanehisa, M., Furumichi, M., Sato, Y., Kawashima, M. and Ishiguro-Watanabe, M. (2023). KEGG for taxonomy-based analysis of pathways and genomes. Nucleic Acids Res. 51, D587–D592.

43. Hagino K, Onuma R, Kawachi M, Horiguchi T. (2013). Discovery of an endosymbiotic nitrogen-fixing cyanobacterium UCYN-A in Braarudosphaera bigelowii (Prymnesiophyceae). PLoS One. 8(12):e81749. doi: 10.1371/journal.pone.0081749. PMID: 24324722; PMCID: PMC3852252.

44. DeYoe, H.R., Lowe, R.L., Marks, J.C. (1992). Effects of nitrogen and phosphorous on the endosymbiont load of Rhopalodia gibba and Epithemia turgida (Bacillariophyceae). J Phycol 28: 773–777.

45. Dutkiewicz, S., B. A. Ward, F. Monteiro, and M. J. Follows (2012), Interconnection of nitrogen fixers and iron in the Pacific Ocean: Theory and numerical simulations, Global Biogeochem. Cycles, 26, GB1012, doi:10.1029/2011GB004039.

46. Herrero, A., Muro-Pastor, A. M., & Flores, E. (2001). Nitrogen control in cyanobacteria. Journal of bacteriology, 183(2), 411–425. 10.1128/JB.183.2.411-425.2001

47. Paerl H. (2017). The cyanobacterial nitrogen fixation paradox in natural waters. F1000Research, 6, 244. 10.12688/f1000research.10603.1

48. Finlay, J. C., Hood, J. M., Limm, M. P., Power, M. E., Schade, J. D., Welter, J. R. (2011). Light -mediated thresholds in stream water nutrient composition in a river network. Ecology, 92, 140–150.

49. Xia, X., S. Zhang, S. Li, L. Zhang, G. Wang, L. Zhang, J. Wang, Li, Z. (2018). The cycle of nitrogen in river systems: Sources, transformation, and flux. Environmental Science: Processes & Impacts 20 (6):863–91. doi: 10.1039/c8em00042e.

50. Lis, H., Shaked, Y., Kranzler, C., Keren, N., & Morel, F. M. (2015). Iron bioavailability to phytoplankton: an empirical approach. The ISME journal, 9(4), 1003–1013. 10.1038/ismej.2014.199

51. Lau, C. K., Krewulak, K. D., & Vogel, H. J. (2016). Bacterial ferrous iron transport: the Feo system. FEMS microbiology reviews, 40(2), 273–298. 10.1093/femsre/fuv049

52. Larson, C. A., Mirza, B., Rodrigues, J. L. M., & Passy, S. I. (2018). Iron limitation effects on nitrogen-fixing organisms with possible implications for cyanobacterial blooms. FEMS microbiology ecology, 94(5), 10.1093/femsec/fiy046. https://doi.org/10.1093/femsec/fiy046

53. Dengg, M., Stirling, C. H., Safi, K., Lehto, N. J., Wood, S. A., Seyitmuhammedov, K., Reid, M. R., & Verburg, P. (2023). Bioavailable iron concentrations regulate phytoplankton growth and bloom formation in low-nutrient lakes. The Science of the total environment, 902, 166399. 10.1016/j.scitotenv.2023.166399

54. Vigneron, A., Cruaud, P., Culley, A. I., Couture, R. M., Lovejoy, C., & Vincent, W. F. (2021). Genomic evidence for sulfur intermediates as new biogeochemical hubs in a model aquatic microbial ecosystem. Microbiome, 9(1), 46. 10.1186/s40168-021-00999-x

55. Bowles, M.W., Mogollón, J.M., Kasten, S., Zabel, M., Hinrichs, K.U. (2014). Global rates of marine sulfate reduction and implications for sub-sea-floor metabolic activities. Science. 344(6186):889–91. doi: 10.1126/science.1249213. Epub 2014 May 8. PMID: 24812207.

56. Arata, P. X., Alberghina, J., Confalonieri, V., Marïa I. Errea, Estevez, J. M., & Ciancia, M. (2017). Sulfated Polysaccharides in the Freshwater Green Macroalga Cladophora surera Not Linked to Salinity Adaptation. Frontiers, 8. 10.3389/fpls.2017.01927

57. Jasem, M. K., Merai, A. A., & Nizam, A. A. (2023). Characterization and in vitro antibacterial activity of sulfated polysaccharides from freshwater alga Cladophora crispata. Access microbiology, 5(7), acmi000537.v5. 10.1099/acmi.0.000537.v5

58. Orellana, L.H., Francis, T.B., Ferraro, M., Hehemann, JH., Fuchs, B.M., Amann, R.I. (2022). Verrucomicrobiota are specialist consumers of sulfated methyl pentoses during diatom blooms. ISME J 16, 630–641, 10.1038/s41396-021-01105-7

59. Klimek, D., Herold, M. & Calusinska, M. (2024). Comparative genomic analysis of Planctomycetota potential for polysaccharide degradation identifies biotechnologically relevant microbes. BMC Genomics 25, doi: 10.1186/s12864-024-10413-z

60. Olapade, O.A., Depas, M.M., Jensen, E.T., McLellan, S.L. (2006). Microbial Communities and Fecal Indicator Bacteria Associated with Cladophora Mats on Beach Sites along Lake Michigan Shores. Applied and environmental microbiology, 89, 10.1128/AEM.72.3.1932-1938.2006

61. Wang, Y., Zhou, P., Zhou, W., Huang, S., Peng, C., Li, D., & Li, G. (2023). Network Analysis Indicates Microbial Assemblage Differences in Life Stages of Cladophora. Applied and environmental microbiology, 89(3), e0211222. 10.1128/aem.02112-22

62. Seyedsayamdost, M. R., Case, R. J., Kolter, R., & Clardy, J. (2011). The Jekyll-and-Hyde chemistry of Phaeobacter gallaeciensis. Nature chemistry, 3(4), 331–335. 10.1038/nchem.1002

63. Abate R., Oon Y.L., Oon Y.S., Bi Y., Mi S. W., Song G., Gao Y. (2024). Diverse interactions between bacteria and microalgae: A review for enhancing harmful algal bloom mitigation and biomass processing efficiency, Heliyon 10 (17) e36503.

64. Cirri, E. and Pohnert, G. (2019), Algae-bacteria interactions that balance the planktonic microbiome. New Phytol, 223: 100–106. 10.1111/nph.15765

65. Segev, E., Wyche, T. P., Kim, K. H., Petersen, J., Ellebrandt, C., Vlamakis, H., Barteneva, N., Paulson, J. N., Chai, L., Clardy, J., & Kolter, R. (2016). Dynamic metabolic exchange governs a marine algal-bacterial interaction. eLife, 5, e17473. 10.7554/eLife.17473

66. Power, M.E.,Chandra, S., Gleick, P., Dietrich, W.E. (2024). Anticipating responses to climate change and planning for resilience in California’s freshwater ecosystems, Proc. Natl. Acad. Sci. U.S.A. 121 (32) e2310075121, 10.1073/pnas.2310075121

