## Supplemental Figures 1 and 2 for "Composition, growth, succession, and function in the *Cladophora* microbiome: insights from quantitative Stable Isotope Probing and NanoSIMS imaging"

S1: Statistical tests on changes in the growth rates of epiphytic bacteria with different functions on Cladophora in the Eel River. (Mendocino County, California). Results from Tukey's pairwise comparisons tests. Error bars represent the 95% confidence interval for the difference in mean growth rates between any two functional groups. A) Early stage (green *Cladophora*). B) Middle stage (yellow *Cladophora*). C) Late stage (red *Cladophora*).

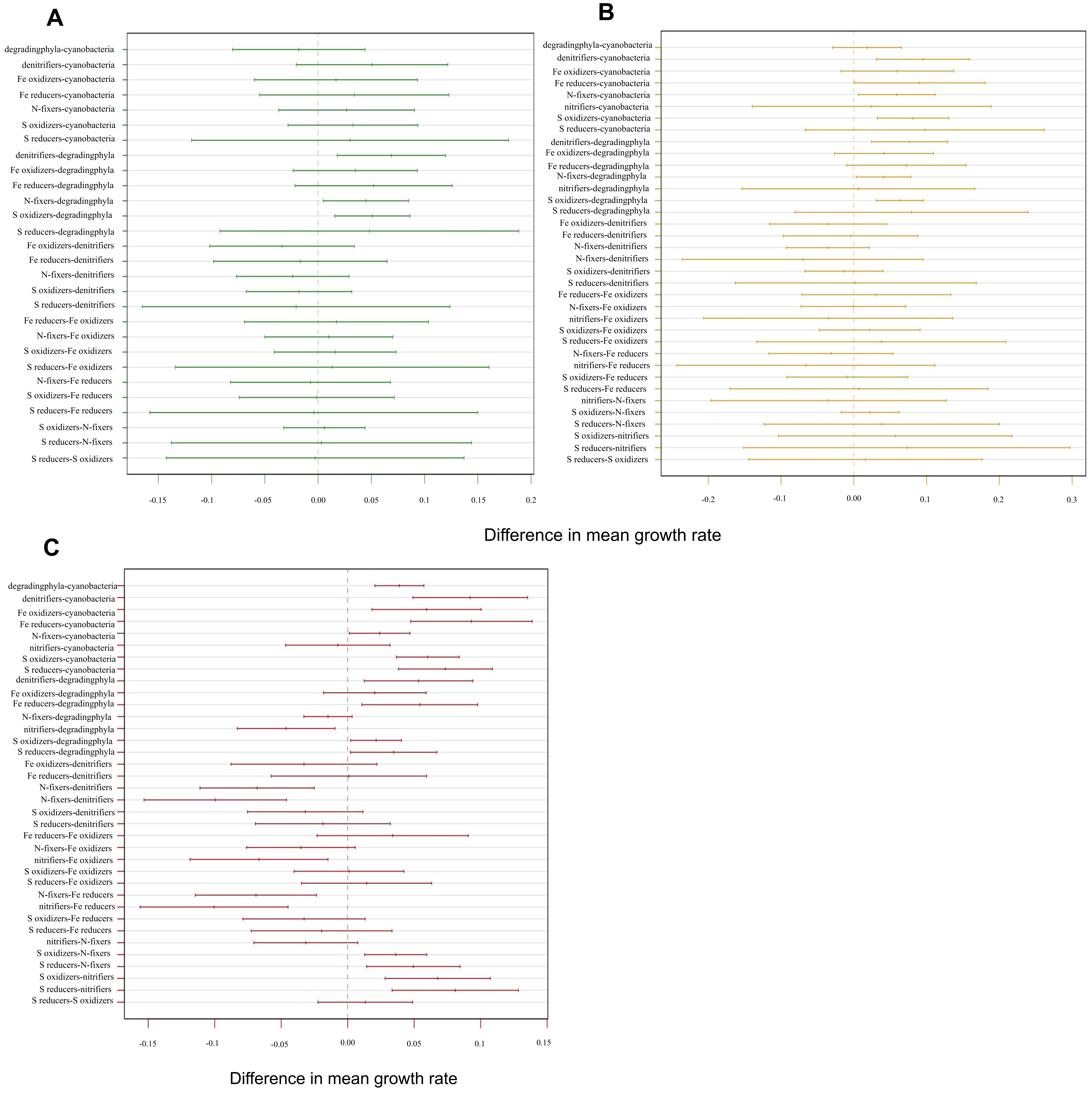

S2: Taxon specific growth rates of bacteria living on Cladophora in the Eel River (Mendocino County, California). Taxa are broken into different functional groups predicted using PICRUSt2 and NCBI.

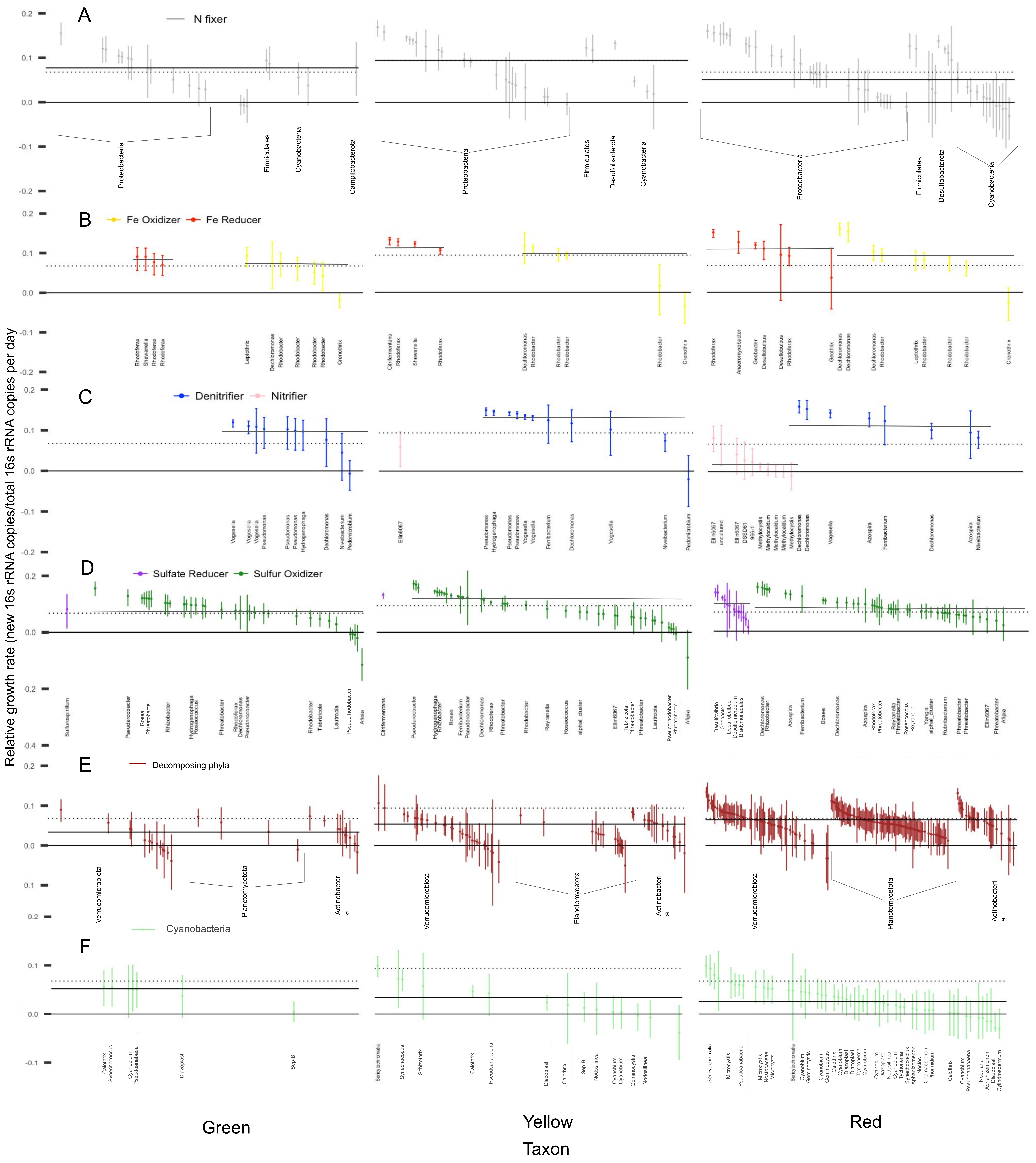
